# Mathematical modeling of genetic pest management through female-specific lethality: Is one locus better than two?

**DOI:** 10.1101/2020.04.06.028738

**Authors:** Michael R. Vella, Fred Gould, Alun L. Lloyd

**Affiliations:** Biomathematics Graduate Program, North Carolina State University, Raleigh, NC 27695, USA; Genetic Engineering and Society Center, North Carolina State University, Raleigh, NC 27695, USA; Department of Entomology and Plant Pathology, North Carolina State University, Raleigh, NC 27695, USA; Department of Mathematics, North Carolina State University, Raleigh, NC 27695, USA

## Abstract

Many novel genetic approaches are under development to combat insect pests. One genetic strategy aims to suppress or locally eliminate a species through large, repeated releases of genetically engineered strains that render female offspring unviable under field conditions. Strains with this female-killing (FK) characteristic have been developed either with all of the molecular components in a single construct or with the components in two constructs inserted at independently assorting loci. Strains with two constructs are typically considered to be only of value as research tools and for producing solely male offspring in rearing factories which are subsequently sterilized by radiation before release. A concern with the two-construct strains is that once released, the two constructs would become separated and therefore non-functional. The only FK strains that have been released in the field without sterilization are single-construct strains. Here, we use a population genetics model with density dependence to evaluate the relative effectiveness of female killing approaches based on single- and two-construct arrangements. We find that, in general, the single-construct arrangement results in slightly faster population suppression, but the two-construct arrangement can eventually cause stronger suppression and cause local elimination with a smaller release size. Based on our results, there is no a priori reason that males carrying two independently segregating constructs need to be sterilized prior to release. In some cases, a fertile release would be more efficient for population suppression.

## Introduction

Insect pests remain a burden to human health and agriculture (World Health Organization 2017; Deutsch et al., 2018). Genetic pest management aims to reduce this burden by releasing engineered insects that either introduce a desired trait into a natural population or reduce the size of the population. There have historically been several large area-wide inundative releases of male insects that were rendered sterile by exposure to radiation (Gould and Schliekelman 2004). In these releases, local elimination of the target species was achieved as females increasingly mated with the sterile males rather than the wild-type males with whom they would produce viable offspring. Instead of using radiation to cause sterility, a contemporary alternative is to genetically engineer strains in which the males cause all their offspring or exclusively their daughters to die or to have low fitness (Alphey 2002). Genetically engineered strains in a number of species have been cage- or field-tested (Wise de Valdez et al., 2011; Ant et al., 2012; Harris et al., 2012; Lacroix et al., 2012; Leftwich et al., 2014; Carvalho et al., 2015; Harvey-Samuel et al., 2015; Gorman et al., 2016).

One approach to developing these functionally-sterile strains involves inserting a repressible, dominant lethal trait, which can be active in both sexes or in females only (Heinrich & Scott, 2000; Thomas et al., 2000). Intuitively, modeling studies have found that female-killing (FK, also sometimes referred to as fsRIDL, or <u>f</u>emale-<u>s</u>pecific <u>r</u>elease of insects carrying <u>d</u>ominant lethals) can be advantageous over bisex lethality (BS) because it kills females while allowing the transgene to propagate through multiple generations in heterozygote males (Schliekelman & Gould, 2000; Thomas et al., 2000). This would seem especially useful when females but not males transmit pathogens. However, heterozygous males can also serve as a reservoir for wild-type alleles, which can make FK less effective than BS under some conditions (Foster et al., 1988; Gentile et al., 2015). It should be noted that BS strains for mosquito disease vectors typically require sex-sorting because release of females would be considered unacceptable. It can also be advantageous to release only males as females do not contribute to genetic suppression and tend to mate with the released males and thus reduce their efficiency (Rendón et al., 2004) except in some situations where there is age structuring in the population (Huang et al., 2009).

For either the BS or FK approach, in order to rear the transgenic strain in the generations prior to release, it must be possible to inactivate the dominant lethal gene. Often this is achieved through a Tet-off system where tetracycline in the diet represses the activator for a lethal gene (Gossen & Bujard, 1992). For an FK strain, the release generation is reared on a diet not containing tetracycline. This results in only males surviving. Further, as the offspring of released FK or BS males would feed on a tetracycline-free diet under field conditions, the lethal gene is turned on and death ensues.

The full molecular design involves two molecular components: 1. the tetracycline-repressible transactivator (tTA) with a promoter, and 2. a lethal gene with an enhancer/promoter consisting of multiple tTA binding sites (tetO) and a core promoter. In the initial two-component systems, tTA was expressed in females by using a female-specific promoter (Heinrich & Scott, 2000; Thomas et al., 2000). The second component was a lethal gene (e.g., proapoptotic) driven by a tetO enhancer-promoter. The two molecular components were built in separate constructs that were inserted independently. Subsequently, a simpler, two-component system was developed in which tTA acts as both the activator and lethal gene. Here, a single construct includes a tTA coding sequence driven by a tetO enhancer-promoter. In this autoregulated system, high levels of the tTA activator cause lethality in late stage larvae or in pupae. The mortality is possibly due to a general interference in transcription (Gong et al., 2005). FK single-construct strains have included a sex-specifically spliced intron from the *transformer* or *doublesex* genes inserted within the tTA gene (Fu et al., 2007). In these FK strains, only the female tTA transcript encodes a functional protein. A different, single-construct approach for FK with *Aedes aegypti* and *Aedes albopictus* uses a female-specific indirect flight muscle promoter from the Actin-4 gene (Fu et al., 2010; Labbé et al., 2012). All field trials with transgenic FK or BS strains have been with single-construct strains.

More recently, two-construct FK strains have been made with an early embryo promoter driving tTA expression and a tTA-regulated lethal gene that contains a sex-specifically spliced intron (Yan et al., 2019). An advantage of these strains is that female lethality occurs at the embryo or early larval stages, which produces considerable savings in larval diet costs in a mass rearing facility. Although it should be possible to develop any two-component systems as a single construct (Yan & Scott, 2015), they are typically developed as independently-segregating constructs. This is because it can be advantageous to insert each component independently into different insect strains, identify the most effective strains, and then produce individuals bearing both components by crossing. The final transgenic insects have the two components located at two, separate loci (Schetelig & Handler, 2012; Ogaugwu et al., 2013; Scott, 2014; Schetelig et al., 2016; Yan et al., 2019).

FK strains with two constructs are generally thought of as useful research tools with potential to be used in rearing facilities so that the final generation before release would only produce males (e.g. Schetelig & Handler, 2012; Ogaugwu et al., 2013; Yan & Scott, 2015). It has been suggested that independent inheritance of the components would cause a breakdown in the female killing in the second generation after release (Ogaugwu et al., 2013; Yan & Scott, 2015). However, previous theoretical studies of FK systems have only modeled the components as being inserted together on a single locus (Thomas et al., 2000; Schliekelman & Gould, 2000; Alphey et al., 2011; Gentile et al., 2015).

Here we evaluate the effectiveness of 1 and 2-locus FK, along with BS for comparison. We use a computational model parameterized for the *Aedes aegypti* mosquito that is a vector for several human pathogens. We explore the release of strains with killing in either juveniles or adults. We show that under reasonable assumptions about fitness costs of the insertions, there is not a substantial difference between the 1 and 2-locus FK approaches, particularly when compared to the differences between FK and BS. These results demonstrate the release potential of recently developed 2-locus FK constructs.

## Methods

Our computational model implements the genetics of FK and BS by separately tracking the number of individuals in the population of each genotype, with genotype denoted by subscript *i*. For the single-locus system (Table 1), we let the transgenic allele be represented by K and the wild-type allele at that locus be represented by k, with a total of *N=3* possible genotypes. For the 2-locus system (Table 2), we let A and B represent the transgenic alleles (i.e., tTA and lethal gene) inserted at two separate loci with wild-type alleles a and b, respectively, for a total of *N=9* possible diploid genotypes.

**Table 1:** 1-locus genotypes, with associated viabilities and fitnesses. Viability of genotype *i*, 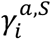, takes the value listed when the approach causes loss of viability in sex *S* (for female killing, only when *S = F*; for bisex, when *S = F* or *S = M*) with timing *a* (for early approaches, when *a = E*; for late approaches, when *a = L*), and are 1 otherwise. Fitnesses 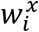 apply for both hatching (*x* = H) and male mating competitiveness (*x* = M).

| $i$ | genotype | $\gamma_i^{a,S}$ | $w_i^x$ |
| --- | --- | --- | --- |
| 1 | kk | 1 | 1 |
| 2 | Kk | 0 | $1 - h \cdot s^x$ |
| 3 | KK | 0 | $1 - s^x$ |

**Table 2:** 2-locus genotypes, with associated viabilities and fitnesses. Viability of genotype *i*, 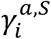, takes the value listed when the approach causes loss of viability in sex *S* (for female killing, only when *S = F*; for bisex, when *S = F* or *S = M*) with timing *a* (for early approaches, when *a = E*; for late approaches, when *a = L*), and are 1 otherwise. Fitnesses 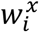 apply for both hatching (*x* = H) and male mating competitiveness (*x* = M).

| $i$ | genotype | $\gamma_i^{a,S}$ | $w_i^x$ |
| --- | --- | --- | --- |
| 1 | aabb | 1 | 1 |
| 2 | aaBb | 1 | $1 - h \cdot s^x(1 - c_A)$ |
| 3 | aaBB | 1 | $1 - s^x(1 - c_A)$ |
| 4 | Aabb | 1 | $1 - h \cdot s^x c_A$ |
| 5 | AaBb | 0 | $1 - h \cdot s^x$ |
| 6 | AaBB | 0 | $1 - h \cdot s^x c_A - s^x(1 - c_A)$ |
| 7 | AAbb | 1 | $1 - s^x c_A$ |
| 8 | AABb | 0 | $1 - s^x c_A - h \cdot s^x(1 - c_A)$ |
| 9 | AABB | 0 | $1 - s^x$ |

We assume complete effectiveness of the constructs, so when there is no gene repression via tetracycline, all individuals bearing the functional BS system and all females with the functional FK system die (i.e. genotype viability of zero). One copy of K is assumed to be sufficient to induce lethality in the 1-locus system, and only one copy each of A and B is required in the 2-locus system. We consider lethality acting at different points in the lifecycle. In insects that experience strong resource competition during larval stages, having the transgene-induced mortality occur during or shortly after the pupal stage, instead of during the egg or larval stages, can yield stronger population suppression. This is because the transgenic juveniles consume resources and thereby increase wild-type juvenile mortality. We model early mortality (E) as occurring in the embryo and late mortality (L) as occurring in pupal stages or in adults before mating, and we assume these differentiate whether the individual contributes toward density-dependent mortality of all individuals in the population. We let 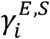 and 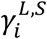 represent the early (embryonic) and late (adult) expected viabilities for individuals of sex *S* and genotype *i*. Tables 1 and 2 give expected viabilities for individuals with each construct and genotype.

We classify constructs into four different approaches depending on when the dominant lethal gene is active as in Gentile et al. (2015): early bisex (EBS), late bisex (LBS), early female-killing (EFK), and late female-killing (LFK). We assume male transgenic homozygotes are released, so mating with wild-type females will produce offspring that are entirely heterozygous, with a copy of each transgene. If the construct(s) affects both sexes (BS), none of these offspring will survive to mate and pass on their genes, making bisex 1-locus and 2-locus equivalent in terms of both population genetics and population dynamics. Female-specific approaches (FK) allow males to continue to propagate the transgenes, and thus inheritance differs between 1-locus and 2-locus approaches. In all, we consider the following six approaches: EBS, LBS, 1-locus EFK (EFK 1), 2-locus EFK (EFK 2), 1-locus LFK (LFK 1), and 2-locus LFK (LFK 2).

Separate from the transgenic, toxin-induced lethality, we account for potential fitness costs caused by the genetic insertion itself. We allow the fitness costs of inserting a novel genetic element to manifest at an early stage as a reduction in the ability of a zygote to survive beyond the egg stage, i.e., the fraction of eggs of that genotype which survive and hatch into larvae. We let the genotype’s hatching fitness, 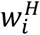, equal the probability of successfully entering the larval stage, with wild-type hatching fitness 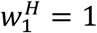. We also allow for transgenic fitness costs to males in the form of reduced mating competitiveness, 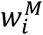, as defined below, with wild-type mating competitiveness 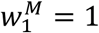.

We generally assume that the fitness costs are equal for the homozygotes in the 1-locus and 2-locus systems to facilitate a direct comparison between the two systems. The 2-locus system requires two separate genetic insertions but is expected to be easier to engineer, which makes equal fitness costs a reasonable base assumption for the purposes of this work (this assumption is relaxed in Figure 3 and Figure S1). We let *s*^*H*^ and *s*^*M*^ be the hatching and mating competitiveness fitness costs, respectively, to the homozygotes KK and AABB, and we allow the two types of costs to vary independently. For simplicity, we assume the degree of dominance for the fitness costs, *h*, is equal for hatching and mating competitiveness. Unless otherwise noted, we assume costs are additive, with *h* = 0.5, such that each copy of the K allele alone contributes a fitness cost of 0.5*s*^*x*^ for the 1-locus system (*x* here indicates that the fitness cost can either be hatching or mating). For the 2-locus system, we allow for unequal fitness costs between each of the insertions. We let two copies of the A allele contribute a fitness cost of *s*^*x*^ *c*_*A*_, where *c*_*A*_ is the proportion of the total 2-locus fitness cost accounted for by the A allele, and one copy of the A allele contributes a fitness cost of *h* · *s*^*x*^ *c*_*A*_. A single B allele contributes a fitness cost of *h s*^*x*^(1 − *c*_*A*_), while being homozygous for B induces a cost of *s*^*x*^(1 − *c*_*A*_). Resulting fitness expressions for all genotypes are listed in Tables 1 and 2.

**Figure 1.**
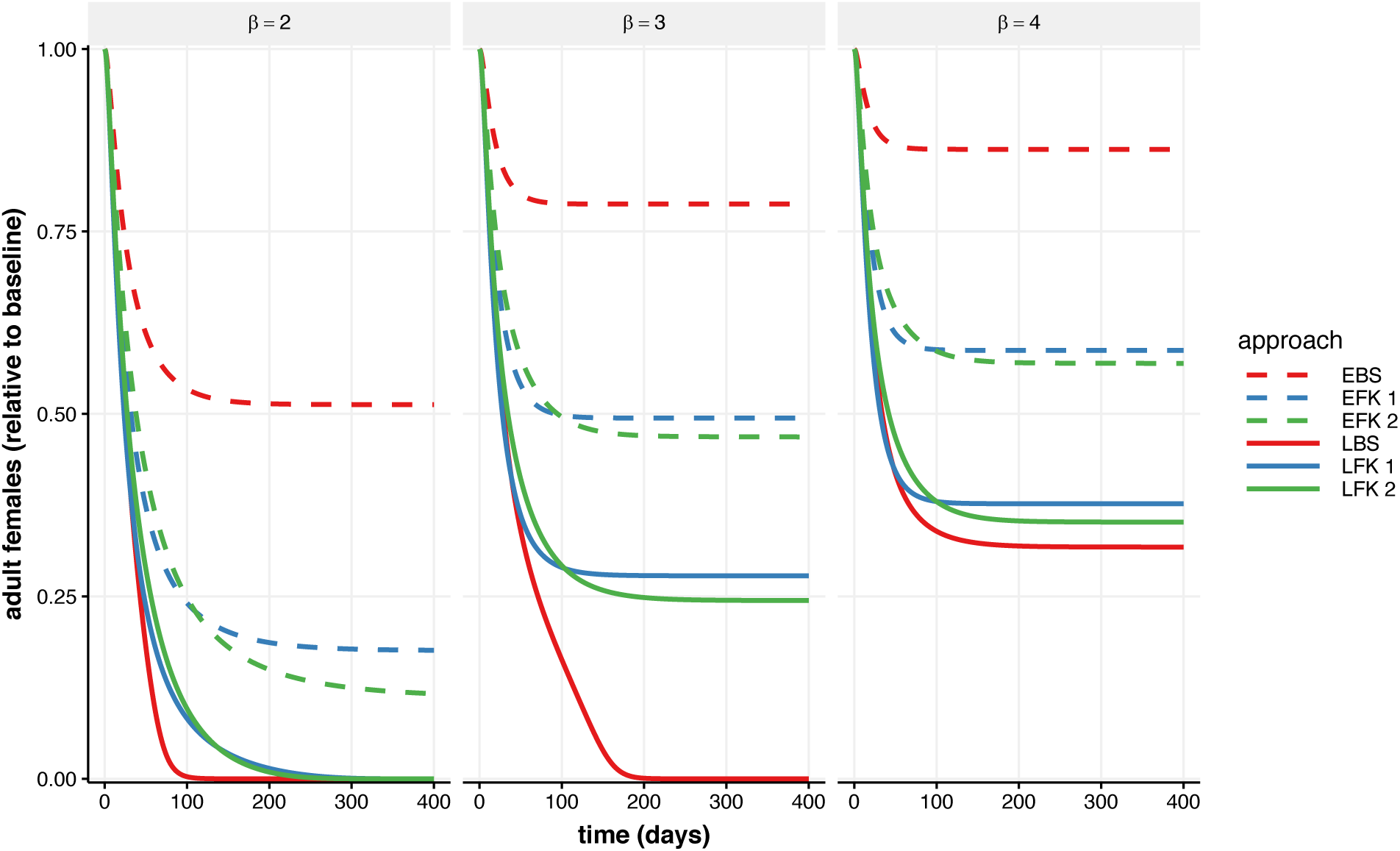
Effect of transgenic releases on population size over time for various strengths of density dependence. The number of viable adult females (relative to pre-release equilibrium) over time is plotted for deterministic simulations with adults for each genetic approach released at a continual weekly release ratio of 1:1 transgenic males to the pre-release equilibrium wild-type males (*r* = 1). Line type differentiates embryo (early-acting, dashed line) and adult (late-acting, solid line) mortality, and line color differentiates bisex (red), 1-locus female-killing (blue), and 2-locus female-killing (green) constructs. Releases are less effective as strength of density dependence (β) increases across panels from left to right. Simulations use fitness parameters *s*^*H*^ *=* 0.2, *s*^*M*^ = 0.1, and *c*_*A*_ = 0.55 and the remaining parameters as listed in Table 3.

**Figure 2.**
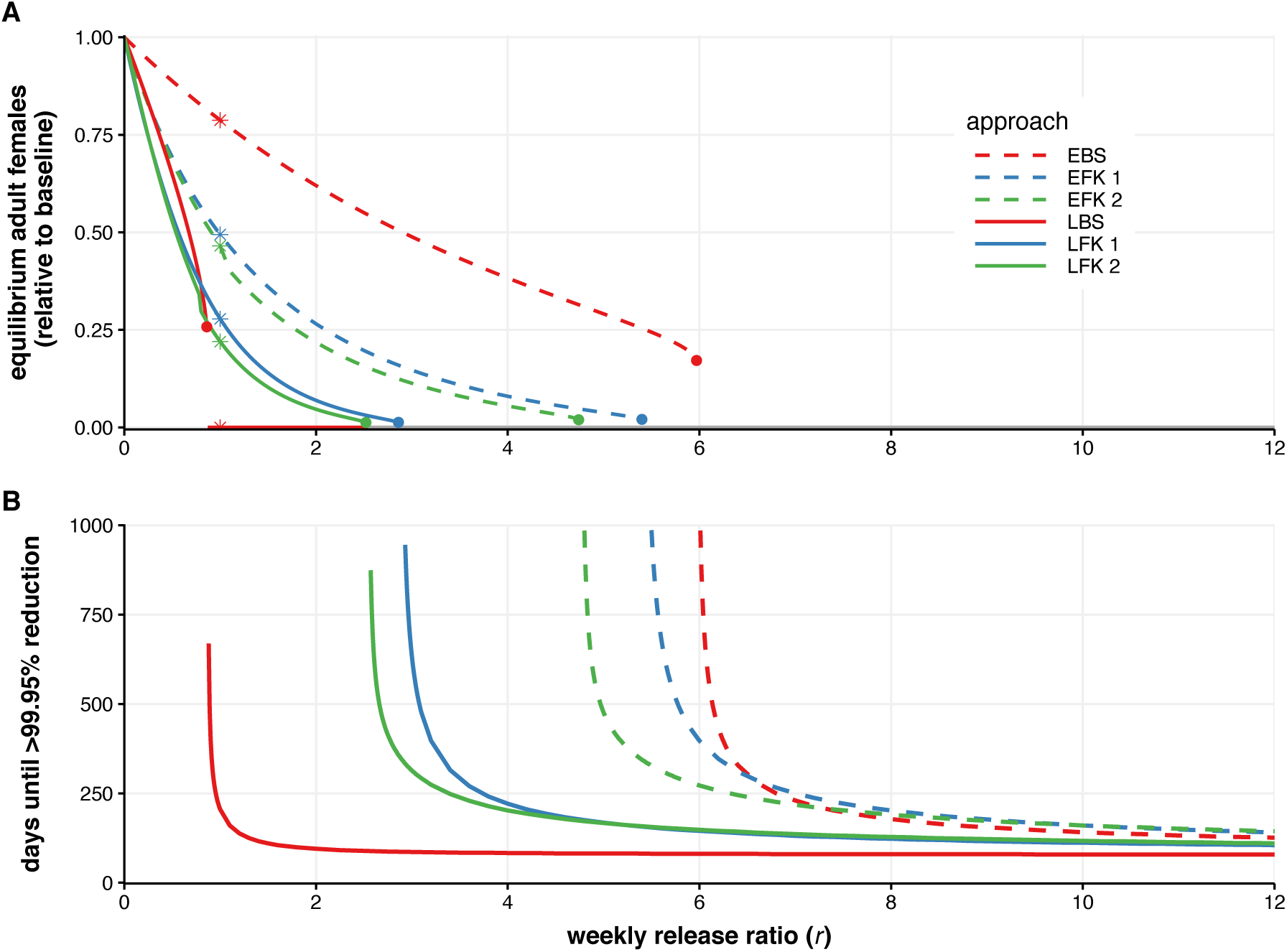
Release outcomes across different release ratios. **A:** Long-term, stable equilibria for number of viable adult females (relative to pre-release equilibrium) for different *r*, found by simulating the system of differential equations until at steady-state. The asterisks indicate *r* = 1, for which the equilibrium values correspond to the middle panel of Figure 1. Each genetic approach exhibits a critical release ratio, *r*_*c*_, above which extinction is guaranteed. Above *r* = 2.52, two or more approaches lead to extinction of the population and hence have equilibria at zero: this is indicated using a grey line. **B:** Time until the number of viable adult females is under 0.05% of equilibrium in deterministic simulations for different *r*. Color and line type match that of Figure 1. Parameters are β = 3, *s*^*H*^ *=* 0.2, *s*^*M*^ = 0.1, and *c*_*A*_ = 0.55, as in Figure 1.

**Figure 3:**
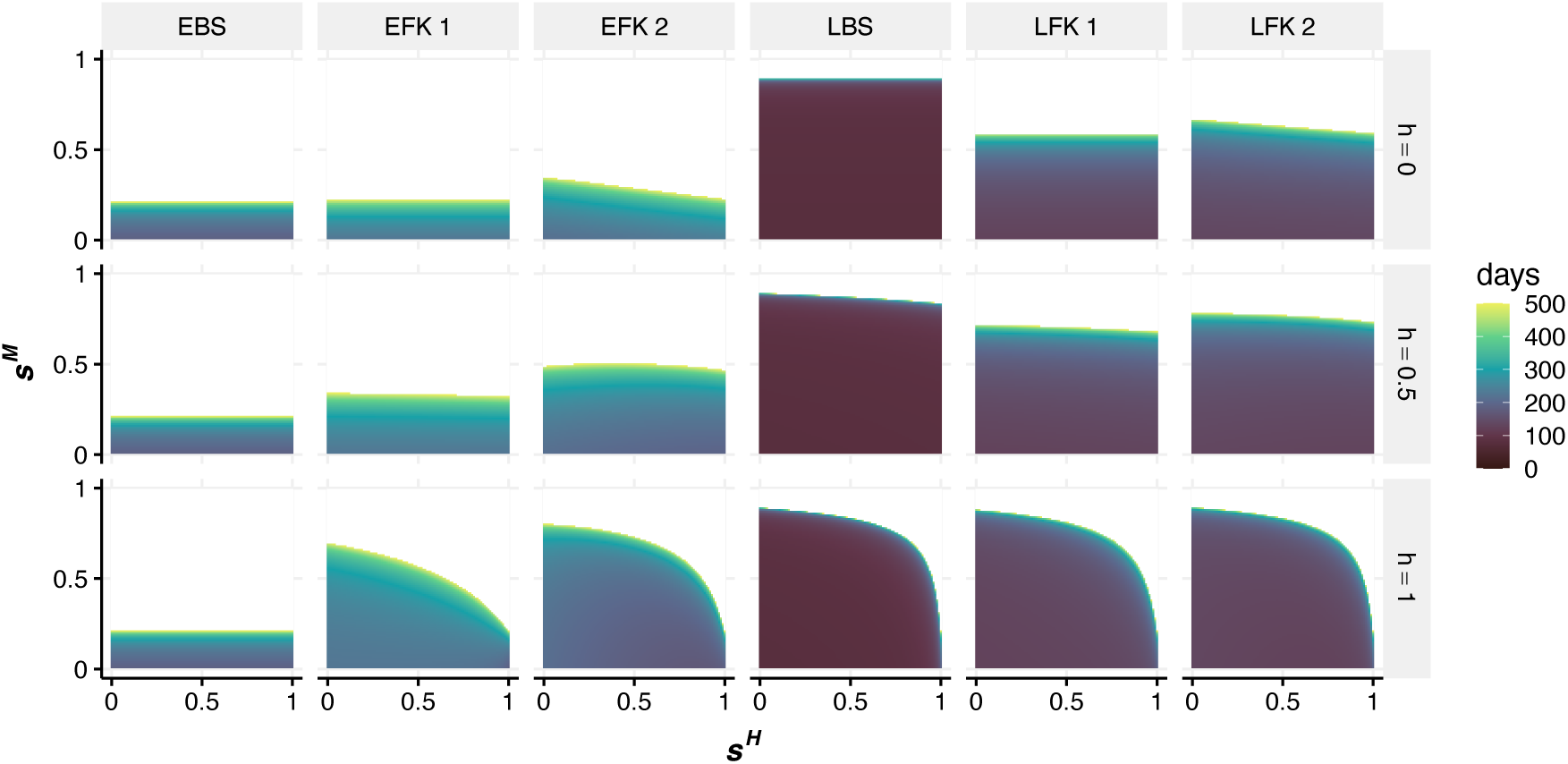
Effects of fitness cost variation on release efficacy for weekly release ratio *r* = 7 and β = 3. Each column of panels shows the results for a different genetic approach, while each row of panels depicts a different degree of dominance, *h*. Within each individual panel, the hatching fitness cost, *s*^*H*^, increases from 0 to 1 along the x-axis, and the cost to male mating competitiveness, *s*^*M*^, increases from 0 to 1 along the y-axis. For every point, a deterministic simulation was run with a unique combination of genetic approach and fitness parameters, and color indicates the number of days until the number of viable adult females is under 0.05% of equilibrium. Darker colors show faster times, with a minimum time of 73 days, and lighter colors show slower times up to 500 days. White areas indicate that the number of females did not fall below the threshold within 500 days. The colored points in the middle row correspond to the times in Figure 2B at *r* = 7.

We model genotype counts over time using a system of ordinary differential equations adapted from Robert et al. (2013). We let 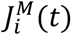 and 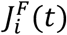 be the number of juvenile (larvae and pupae) males and females, respectively, of genotype *i* at time *t*, and 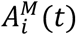 and 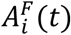 be the number of viable adult male and adult female mosquitoes in the population, respectively, of genotype *i* at time *t*. This gives a maximum of 12 classes of individuals to track (each with different combinations of the 3 genotypes, 2 sexes, and 2 age classes) for the 1-locus system and 36 classes for the 2-locus system, though lethality from the genetic construct prevents survival of certain classes. For instance, EBS only has five non-zero classes (wild-type male and female juveniles and adults, and male adult homozygote transgenic, which are released). We also assume a 50:50 sex ratio and equal hatching fitness costs between males and females, allowing a further reduction in the number of unique classes for late acting approaches because 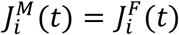 for all *i* and t. This results in 7 classes for LFK 1 (after removing 3 juvenile classes and 2 non-viable adult female classes) and 23 classes for LFK 2 (after removing 9 juvenile classes and 4 non-viable adult female classes). These dimensionality reductions can be useful when finding analytical solutions, but for simplicity, we computationally simulated all 12 (1-locus) or 36 (2-locus) equations.

Accounting for fitness costs, adult females produce juveniles of genotype *i* at time *t* at rate

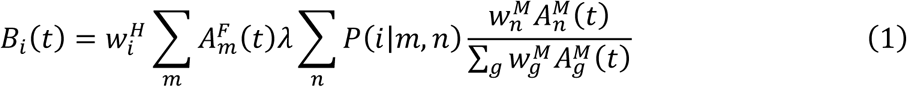

where *λ* is the per-capita birth rate and *P*(*i*|*m, n*) is the probability that a juvenile produced from a mating between a female and male of genotypes *m* and *n*, respectively, will be of genotype *i*. The fraction gives the probability that a randomly chosen male adult is of genotype *n*, weighted by mating competitiveness 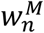. The offspring genotype probabilities are calculated assuming Mendelian inheritance, and for the 2-locus case, independent segregation of genes at each locus. Because of the 50:50 sex ratio, half of the hatching juveniles are male and half are female.

Juveniles of each genotype and sex emerge to adulthood at per-capita rate *v*. We assume juveniles, adult males, and adult females have per-capita density-independent mortality rates of *μ*_*s*_, *μ*_*M*_, and *μ*_*C*_, respectively. Juveniles also undergo density-dependent mortality at a per-capita rate (*αJ*)^*β*−1^, where *J* is the total number of juveniles. The strength of density dependence is adjusted by varying β, with higher β resulting in a faster return to equilibrium population size after a small perturbation. A value of β = 2 gives the logistic model for population dynamics. By default, we let β = 3 to model an environment which would be more difficult for successful suppression (e.g., Hibbard et al., 2010). The equilibrium size of an entirely wild-type population varies with α, and to keep simulations with different values of β comparable, we choose the value of α so that the equilibrium number of wild-type females remains the same.

We assume a continuous release of homozygote engineered males at (daily) rate 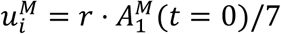, where *r* is the weekly release ratio (engineered:wild-type) based on the equilibrium number of males prior to the release. By maintaining a constant number of released males, the effective release ratio increases as the population size decreases. The release genotype is KK for 1-locus (*i* = 3) and AABB for 2-locus (*i* = *H*), and because we assume that no females are released, 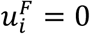 for all *i*.

The resulting system of differential equations (with time dependence of 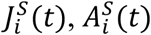, and *B*_*i*_(*t*) omitted for simplicity of notation) is

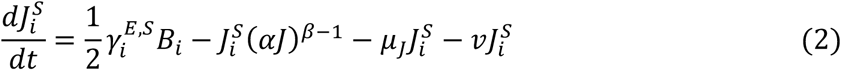

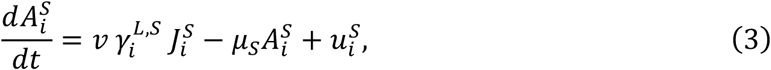

for *i* =1..*N* and *S* = F or M. All model parameters are listed in Table 3 and are based on those used for *Ae. aegypti* by Robert et al. (2013). All numerical simulations of the differential equations have initial conditions at the wild-type equilibrium.

**Table 3:** Model parameters.

| Parameter | Description | Value/range | Reference |
| --- | --- | --- | --- |
| $\mu_J$ | Density-independent per-capita juvenile mortality rate | 0.03 day<sup>-1</sup> | Rueda et al. 1990 |
| $\mu_M$ | Adult male per-capita mortality rate | 0.28 day<sup>-1</sup> | Muir and Kay 1998, Fouque et al. 2006 |
| $\mu_F$ | Adult female per-capita mortality rate | 0.10 day<sup>-1</sup> | Muir and Kay 1998, Fouque et al. 2006 |
| $\lambda$ | Female per-capita larval production rate | 8 day<sup>-1</sup> | Harrington et al. 2001, Styer et al. 2007 |
| $v$ | Per-capita rate of emergence to adulthood | 0.14 day<sup>-1</sup> | Muir and Kay 1998 |
| $\beta$ | Strength of density dependence | 2 to 4 | |
| $A_1^F(0)$ | Equilibrium number of wild-type females | 2000 | |
| $r$ | Weekly release ratio of transgenic: equilibrium wild-type males | 0 to 12 | |
| $\alpha$ | Density dependence parameter chosen based on $\beta$ and $A_1^F(0)$ | | |
| $w_i^H, w_i^M$ | Egg hatching and male mating competitiveness fitnesses of genotype $i$ | See Tables 1 and 2 | |
| $s^H, s^M$ | Egg hatching and male mating competitiveness fitness costs to homozygotes | 0 to 1 (0.2 and 0.1 default, respectively) | |
| $h$ | Fitness cost degree of dominance (percent of $s^x$ incurred in heterozygotes) | 0 to 1 (0.5 default) | |
| $c_A$ | Proportion of $s$ contributed by A (2-locus) | 0.5 to 1 (0.55 default) | |
| $\gamma_i^{E,S}, \gamma_i^{L,S}$ | Viabilities of genotype $i$ for early (E; embryonic) and late (L; adult) approaches in sex $S$ | See Tables 1 and 2 | |

In order to explore the effects of demographic stochasticity and genetic drift, we also run simulations using an analogous continuous time Markov chain model (for details, see Appendix 1).

## Results

Each of the FK and BS genetic strategies has the goal of causing the population to decline by reducing the number of reproductive adult females. The strength of density-dependent mortality moderates the reductions in population size because stronger density dependence (higher β) causes the juvenile mortality rates to decrease more quickly as population size decreases from equilibrium. In a system with strong density dependence, the weekly release ratio (*r*) must be larger to achieve the same amount of population suppression as in a system with weak density dependence. Large *r* can result in target population extinction. In contrast, small *r* results in a new, lower equilibrium population density, where the proportion of individuals that die due to bearing the transgene is not high enough to outweigh the increased survival of juveniles due to decreased density-dependent mortality in the smaller population.

Figure 1 demonstrates the outcome of release for each genetic approach at *r* = 1 under the deterministic model (see Figure S1 for time series with multiple release ratios). At this release ratio, approaches that are late-acting (i.e., mortality in pre-mated adults, shown by solid lines) reduce the number of viable females to a lower number than approaches that are early-acting (i.e., mortality in the embryonic life stage, shown by dashed lines). BS approaches (red) reduce the number of females faster and to lower levels than female-specific approaches (blue/green) with late-acting mortality, but the opposite is true for early-acting mortality. Among FK approaches, 1-locus reduces the number of females more quickly initially, but 2-locus eventually suppresses the population slightly more than 1-locus. Overall, this suggests that LBS is most effective, followed by LFK, EFK, and EBS, with little difference between 1 and 2-locus FK. Under stronger density dependence (higher values of β), releases result in weaker suppression of the population, with none of the approaches causing extinction of the population if β = 4.

In general, large release ratios result in extinction (the population goes to an equilibrium size of zero), and small release ratios result in a suppressed but non-zero equilibrium population size. For most sets of parameters, there is a critical release ratio, *r*_*c*_, above which the release is large enough to cause the population to go extinct. With such a large number of released males, population extinction is the only stable equilibrium, meaning the release will cause extinction regardless of initial population size. For ongoing release at release ratios below *r*_*c*_, there is a non-zero stable equilibrium for the number of viable adult females, meaning that release will not push a wild-type population to extinction. A population size of zero is also stable, meaning the release could protect against re-invasion of the wild-type if continued after population extinction, but the system will approach the non-zero equilibrium unless starting from very low population sizes, i.e., it is a bistable system with a low invasion threshold (as shown in Figure S2 with time series starting from multiple initial conditions). Mathematical details on the shift in qualitative behavior at *r*_*c*_ and analysis using Mathematica (Wolfram Research, 2019) can be found in Appendix 2 and Figure S3.

Figure 2A shows how release ratios affect stable equilibria when β = 3, with only the non-zero stable equilibrium plotted when the system is bistable. Each of the lines is discontinuous, jumping from a non-zero equilibrium to a zero equilibrium at *r*_*c*_. The smallest release size required to cause extinction for each approach is 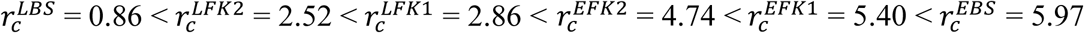. This order of effectiveness matches that of Figure 1 and mirrors the results for single locus constructs in Gentile et al. 2015. However, when *r* is smaller than 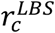 and extinction does not occur, LBS has a higher equilibrium population size than either of the late FK methods. There is a small relative difference between the different FK approaches, with similar equilibrium sizes when *r* is below the critical release ratios and 1-locus FK requiring a release ratio less than 15% larger than 2-locus FK to cause extinction.

In settings where release causes the population to go extinct, we can consider the time it takes to reach extinction after starting release. Figure 2B shows the time it takes for the number of females to reach less than 0.05% of the pre-control equilibrium (suppression to below 1 adult female when starting from a pre-release equilibrium of 2000) in deterministic simulations, which also reflects the time to extinction in stochastic simulations (see Figure S4). Once release ratios are high, LBS drops the population under 0.05% of the equilibrium faster than late FK methods. Also, both of the 1-locus FK approaches are slightly faster than the respective 2-locus FK approaches, though the differences are small for practical purposes. At *r* = 12, for example, the times are 86 (LBS), 114 (LFK 1), 119 (LFK 2), 133 (EBS), 149 (EFK 1), and 153 (EFK 2) days. The results are similar for β = 2 (see Figure S5).

Overall, FK 1 and FK 2 are quite similar. If either of the A or B alleles in 2-locus FK becomes fixed in the population, the 2-locus approach becomes nearly identical to 1-locus FK, where one copy of the unfixed allele causes mortality in females. For example, if all individuals in the population already have the B allele, only one copy of the A component is additionally necessary, just as a single copy of the K allele causes mortality. In this case, the long-term equilibrium can be identical to 1-locus FK, though a fitness cost to the fixed allele decreases the average fitness of the entire population and makes the population size lower for 2-locus FK than 1-locus FK. Whether one of the 2-locus FK alleles become fixed depends on the fitness costs and release ratios of the system (see Appendix 3, Figure S6, and Figure S7).

An observation from Figure 2A is that LBS, LFK 2, and LFK 1 cause a similar level of suppression when the release ratio is near *r* = 0.8. Previous work has suggested that LFK allows the wild-type allele to propagate in heterozygous males, making it less effective than LBS (Gentile et al., 2015). This is true at high release ratios, but not at all release ratios. Without fitness costs, the strategies are equally effective if the number of released males is equal to the number of wild-type males in the population at equilibrium: heterozygotes carry both wildtype and transgenic alleles, and therefore the survival of heterozygous males does not affect allele frequency. At small release ratios, when LBS release results in low transgenic frequency and thus a large equilibrium population size, survival of heterozygous males would allow the transgenic allele to propagate further and increase in frequency, explaining why LFK has a lower equilibrium than LBS in this narrow window of small releases.

The main difference exhibited between FK 1 and FK 2 can also be explained by their propagation of the transgenic and wild-type alleles. When the components are separated across two loci, the A and B alleles become unlinked, with some individuals only inheriting one allele or the other, while having linked components guarantees inheritance of the transgenic allele and reduces the population size more quickly initially. Eventually, however, the accumulation of transgenic alleles in the 2-locus system causes production of a higher proportion of unviable genotypes and greater population suppression (Figure 2).

Results over a wide range of fitness parameters highlights the minimal differences between FK 1 and FK 2. Figure 3 shows the times until 99.95% reduction in viable adult females with a release ratio of *r* = 7, which is larger than *r*_*c*_ for all approaches in the base parameter set, β = 3, and varying sets of fitness parameters. Light colors show parameter sets that resulted in simulations that took longer to reduce the population size, and dark colors show shorter times, while white areas indicate that the population was not reduced by 99.95% within 500 days. Within each panel, which represents a single approach (columns of panels) and degree of dominance, *h*, (rows of panels), there is a similar pattern. At low fitness costs (hatching, *s*^*H*^, varying across the x-axis, and mating competitiveness, *s*^*M*^, varying across the y-axis), the release reduces the population below the threshold, and there is fairly little variation between times (colors) within these regions. As one or both of the fitness costs increase, there is a margin of longer times (light color) separating the successful suppression region from the unsuccessful (white) region. As evident from Figure 2, long times to reduce the population indicate that the release ratio of *r =* 7 is only slightly greater than *r*_*c*_ for those parameter values, and a system with increased fitness costs have *r*_*c*_ > 7, so there is not suppression to 0.05% of equilibrium.

There is little difference between FK 1 and FK 2, particularly for LFK, since the release ratio is much higher than *r*_*c*_ in most of the region with successful release. Even with different fitness costs, FK 1 and FK 2 do not differ drastically in time until extinction; FK 2 would be substantially less effective only if the costs become large enough for the release to become too small to cause extinction (i.e., *r* becomes smaller than *r*_*c*_ because of the higher costs). Unpacking the differences between all approaches, for the EBS approach, neither *h* nor *s*^*H*^ affect the length of time for reduction; only released males experience costs so the degree of dominance does not affect results, and all offspring that inherit a transgene are inviable so hatching costs do not affect results. For EBS, increasing the cost to male mating competitiveness drastically decreases release efficacy because doing so effectively reduces the release ratio (fewer of the released males successfully mate and contribute to population reduction), and with no surviving transgenic offspring, the release ratio is the main contributing factor to EBS efficacy. When *s*^*H*^ = 1 and *h =* 1, none of the offspring survive for the EFK 1 and EFK 2 approaches, making the times shown identical to EBS. Decreasing these parameters makes a difference for FK approaches because the males are subject to fitness costs. Dominant fitness costs (bottom row) actually enable successful population reduction for higher male mating costs. The explanation for this effect relates to the propagation of wild-type alleles, similar to the previous description: homozygote males that are being released are affected regardless of *h*, and when *h* > 0, the fitness cost also reduces the spread of wild-type alleles in heterozygotes. This effect is also evident in other transgenic systems, such as two-locus underdominance, where dominant transgenic allele fitness costs prevent the wild-type from being maintained in the population at small frequencies (Dhole et al. 2018).

## Discussion

The recent literature on FK systems makes the assumption that strains built with constructs inserted at two independent loci will not be as useful for field releases as those built with a single construct. The assumption is that the two constructs will separate from each other in the second generation after a release and will become non-functional. Our modeling results demonstrate that a 2-locus FK (FK 2) should behave similarly to a 1-locus FK (FK 1) and would not present any significant disadvantages in its ability to suppress a population. We generally made the assumption that because of the flexibility of the 2-locus engineering approach, the total fitness cost for the two insertions would be similar to the single insertion at 1-locus. If the total cost of the 2-locus approach was less than for the 1-locus approach, the 2-locus approach would likely be preferred. The reverse also holds. Importantly, based on our results, there is no a priori, general reason for genetic engineers to favor a 1-locus system. The choice will likely depend on specific biological and genetic characteristics of the target species.

Assuming equal costs, FK 1 is slightly faster at initial population reduction, but FK 2 can eventually suppress the population to lower numbers. FK 2 also has a slightly lower critical release ratio than FK 1, meaning a smaller release size is necessary to guarantee extinction. For many combinations of fitness costs and release ratios, one of the FK 2 alleles would be driven to fixation, resulting in a genetic system similar to FK 1. The differences between FK 1 and FK 2 are much smaller than between FK and BS approaches. Comparing FK and BS approaches, our results are generally similar to previous work (Gentile et al., 2015). Late acting approaches cause extinction with a lower release ratio than early acting approaches, with LBS causing extinction with a lower release ratio than LFK, and EFK causing extinction with a lower release ratio than EBS.

While our modeling results indicate that LBS outperforms the other methods, there are other considerations that affect which approach may be best suited for a given scenario. In our model, parameterized for mosquitoes, density-dependent mortality during early life-stages was an important factor and caused early-acting approaches to result in less population reduction than late-acting approaches. In species with little density-dependent dynamics in juveniles, the difference in effectiveness between early and late-acting would be minor, though this is not the case for many pest species.

Beyond population dynamics, there are economic and social factors that differ between approaches. For some systems, it will be necessary to engineer constructs into lab strains and then backcross the construct or constructs into a strain that have a genetic makeup similar to the targeted population. In general, it should be easier to do the backcrossing with a one-locus system. Rearing costs are also expected to vary between approaches. With EFK, juvenile females experience mortality before consuming food, whereas EBS, LBS and LFK require rearing of juveniles of both sexes. Furthermore, BS approaches require sexing to remove females prior to release, which increases the total rearing costs and is often difficult to do with complete accuracy. When releasing a species that is a disease vector, sexing accuracy is meaningful from a social perspective as release of females could contribute to disease transmission.

Apart from engineering and rearing, for most agricultural pests, the juvenile stages of males and females cause damage to crops and livestock. In the first generations of transgenic pest releases, the late acting approaches will leave more feeding immatures in the environment. For LFK and EFK, male immatures will still cause damage. This is unlikely to be favored by farmers even though the overall population could be decreasing rapidly, and EBS could be preferred. Finally, even if late-acting mortality may be ideal for a given scenario, controlling the timing of mortality at the intended life stage may not always be feasible, for example due to leaky expression of the lethal gene.

The model used here has several limitations. An important factor that could affect population genetics is spatial heterogeneity. For example, in a spatial model of FK 2, it would be possible, particularly in small populations, for different patches to have different transgenic alleles reach fixation. A spatial model would also be useful to determine if FK 2 has any differences in resilience to wild-type reinvasion. The details of such a spatial model, including rates of release, would depend on species. A species-specific model could also implement different forms of density-dependence, age-structure, and mating parameters. Finally, given its generality, our model does not account for any potential mechanisms for resistance development. Depending on the mechanism of lethality, there may be advantages for having both components for lethality inserted together. While these areas require further investigation, our results indicate that overall, there is little difference in the pest population suppression efficacies of 1-locus FK and 2-locus FK.

## Acknowledgements

This research was funded by the Research Training Group in Mathematical Biology, funded by NSF grant RTG/DMS-1246991 (MRV and ALL), NSF IGERT grant 1068676 (MRV, FG and ALL), NIH grant 1R01AI139085-01 (FG and ALL), and the NC State Drexel Endowment (ALL). We thank M.J. Scott, S. Dhole, B. Hollingsworth, J. Baltzegar, and J. Sudweeks for comments and helpful discussion.

## Notes

### Competing Interest Statement

The authors have declared no competing interest.

