## Supplemental Information for "Mathematical modeling of genetic pest management through female-specific lethality: Is one locus better than two?"

**Appendix 1: Stochastic simulations**

In order to understand the importance of demographic stochasticity (e.g. to simulate extinction events) and genetic drift, we formulated a stochastic model that is analogous to our deterministic model. Specifically, the rates of each process (birth, maturation and death) that appear in the deterministic model were taken to be rates of a continuous-time, discrete-state Markov process model. This model was simulated using a tau-leaping approach. The steps are as follows: 1. calculate each of the rates $k_j(t)$ in the system at time $t$, 2. advance the time step by the time step $\tau$, 3. approximate the number of times each event occurred using a Poisson distribution with mean $\tau k_j(t)$, and 4. adjust the states accordingly before repeating all steps. We ran simulations using the adaptivetau package in R (Johnson 2014), which uses automatic selection of $\tau$ with adaptive explicit-implicit tau-leaping (Cao et. al. 2007).

For each approach, we ran 300 stochastic simulations for each value in a range of release ratios with the default parameters from the main text. Simulations were stopped after 1000 days. The mean time until there were no adult females in the stochastic simulations (Figure S4) is similar to the time until 99.95% reduction in deterministic simulations (as shown in Figure 2B). In stochastic simulations, random fluctuations can cause the population to go extinct even when $r$ is below $r_c$. As $r$ approaches $r_c$ and the equilibrium population size becomes smaller, extinction occurs within 1000 days in a higher percentage of stochastic simulations. Because the mean is taken from the simulations that go extinct, the mean time until extinction is biased downwards, most notably when extinction times frequently exceed 1000 days. For this reason, Figure S4 only shows outcomes when at least 200 of the 300 simulations reached extinction. While smaller wild-type populations would increase the variance in stochastic simulations, the results suggest the deterministic simulations represent the overall dynamics well.

**Appendix 2: Equilibrium analysis**

For the 1-locus system, we analytically solved for equilibria and calculated their stabilities using the eigenvalues of the Jacobian – a standard analytic approach for dynamical systems (Strogatz, 2001). The system exhibits a saddle-node bifurcation as $r$ increases. Here this means that when $r$ is small, there is a non-zero stable equilibrium for number of viable females, but past a critical release ratio, $r_c$, the only stable equilibrium is a population size of zero. The bifurcation diagram is shown in Figure S3, which shows the same stable equilibria as Figure 2a but also shows unstable equilibria. For the 2-locus system, we could not find an analytical solution, but simulation results indicate similar dynamics are present.

Below $r_c$, the system is bistable, with dynamics bringing the population to extinction if beginning below the unstable equilibrium and bringing the population to a non-zero, stable equilibrium when beginning above the unstable equilibrium. While the unstable equilibrium which serves as the threshold for whether the system goes to zero or goes to the non-zero equilibrium is in multiple dimensions, there is also a threshold when beginning from a wild-type only population. The bistability in a 1-locus LFK system is illustrated in Figure S2. Simulations beginning with low numbers of wild-type individuals (the initial number of juveniles and males are reduced equally to the number of females) go to extinction instead of the non-zero, stable equilibrium. This means that even when the release ratio is below $r_c$ in a system, release could bring the population size to zero following population reduction by another method, such as spraying pesticides. In practice, however, it could be difficult to achieve enough suppression to bring the system below the unstable equilibrium. Additionally, in an area already without any wild-type individuals, ongoing release below $r_c$ could prevent (small-scale) immigration of wild-type from re-establishing a population.

**Appendix 3: 2-locus population genetics with additive fitness costs**

As noted in the main text, if either the A or B alleles are at fixation in a population, the population genetics of 2-locus FK becomes effectively equivalent to 1-locus FK. For example, if B is at fixation, all that is needed is a single copy of A to cause lethality. For some sets of fitness parameters, either A or B is driven to fixation, while other parameters result in intermediate frequencies of both alleles. This effect is illustrated in Figure S6A, which shows allele frequencies over time for both 1-locus and 2-locus LFK with parameters $c_A = 0.75$, $s^M = 0.25$, and $h = 0.5$, and several different values of $s^H$. When $s^H = 0.1$ (left column), the B allele goes to fixation in most simulations, but the A allele can go to fixation in stochastic simulations despite having higher costs than B. The A allele would also go to fixation in deterministic simulations if starting from a much higher frequency than the B allele. At moderate costs (middle column), the B allele always goes to fixation, and at high costs (right column), both alleles reach intermediate frequencies. EFK exhibits similar behavior.

The possible allele outcomes for 2-locus LFK with different combinations of fitness parameters are shown in Figure S7. Here, simulations were conducted with additional transgenic adult male and females in the system at time 0. One set of simulations began with the A allele at 0.5 frequency in adults by adding AAbb adult males and females (in number equal to the wild-type adult equilibria) to the population, while another set similarly began with extra aaBB adults. When fitness costs are equal between the A and B alleles ($c_A = 0.5$, left column), the inheritance of A and B is completely symmetric in deterministic simulations. At high release ratios, the population goes to extinction regardless of initial condition (grey areas), but at lower release ratios, there are two possible outcomes. The A and B alleles are attracted toward equal intermediate frequencies when $s^H$ is high compared to the release ratio (with the intermediate frequency indicated by shades of green). When $s^H$ is small and the release ratio is large enough, the system is attracted toward an equilibrium with whichever of the alleles begins at higher frequency reaching fixation (dark red areas), while the other allele reaches an intermediate frequency. In these cases, the hatching fitness cost is small enough to be outweighed by the influx of AA or BB in released adult males.

When $c_A > 0.5$ (middle and right columns), some parameter sets result in the B allele reaching fixation even in simulations beginning with a higher frequency of A (black areas). As $c_A$ increases, the B allele is pushed to fixation in systems with larger values of $s^H$ because the cost to the B allele is smaller. Likewise, the higher cost of the A allele decreases the (dark red) region where it reaches fixation. In a small region of parameter space, the A allele increases in frequency and imposes a large enough genetic load to cause population extinction, while the B allele would reach fixation without causing extinction (light right areas). As $s^M$ increases, the primary effect is to reduce the effective release size, and thus larger releases are needed to drive an allele to fixation.

In terms of population suppression, the efficacy of 2-locus FK does not significantly depend on the allele frequency dynamics. The amount of suppression is slightly greater when either the

A or B allele reaches fixation as long as there are fitness costs; the costs to the fixed allele impose a genetic load on the entire population. This phenomenon is responsible for sudden, minor decreases in the 2-locus equilibria in Figure 2A and Figure S5A, which happens when the release ratios are large enough to cause one allele to become fixed. However, compared to 1-locus FK, 2-locus FK results in greater suppression even when neither allele is fixed because of the additional propagation of transgenic alleles, as described in the main text. As shown in Figure S6B, the number of adult females over time reaches a smaller number in 2-locus LFK than in 1-locus LFK regardless of whether an allele reaches fixation, with a larger difference in suppression when $s^H$ is larger.

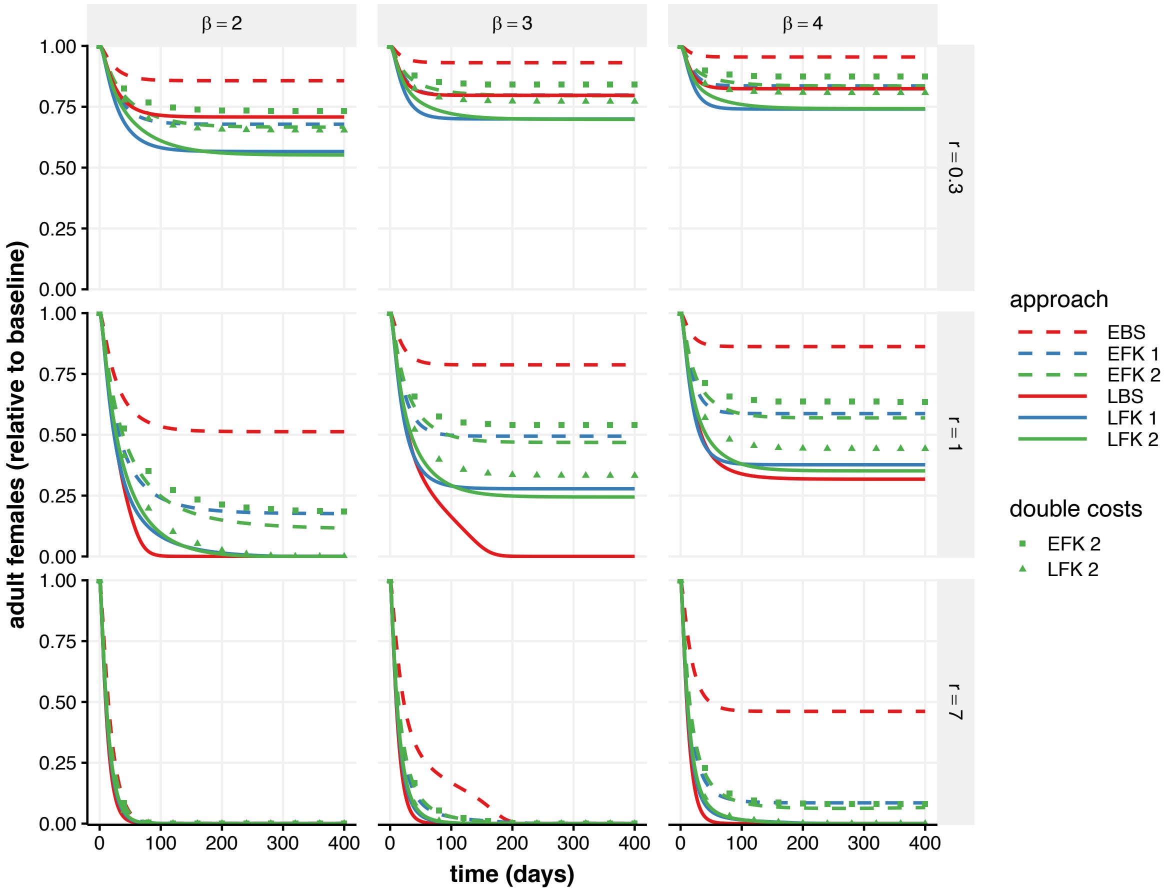


**Figure S1: Effect of transgenic releases on population size over time for various strengths of density dependence and release ratios.** The number of viable adult females (relative to pre-release equilibrium) over time is plotted in deterministic simulations. Release ratios vary across rows, with adults for each genetic approach released at a continual weekly release ratio of 3:10, 1:1, and 7:1 transgenic males to the pre-release equilibrium wild-type males (middle row is equivalent to Figure 1). Line type and colors vary by approach as in the main text, with the addition of squares and triangles to show 2-locus EFK and 2-locus LFK with double the total fitness costs ($s^H = 0.4$, and $s^M = 0.2$, with $c_A = 0.55$). The remaining simulations use the default parameters from the main text: $s^H = 0.2$, $s^M = 0.1$, $c_A = 0.55$, and the remaining parameters as listed in table 3. As strength of density dependence (β) increases and release size (*r*) decreases, the releases are less effective. If 2-locus FK has higher costs (symbols), it can become less effective than 1-locus FK, but at high release ratios, there is little difference in the number of total females over time.

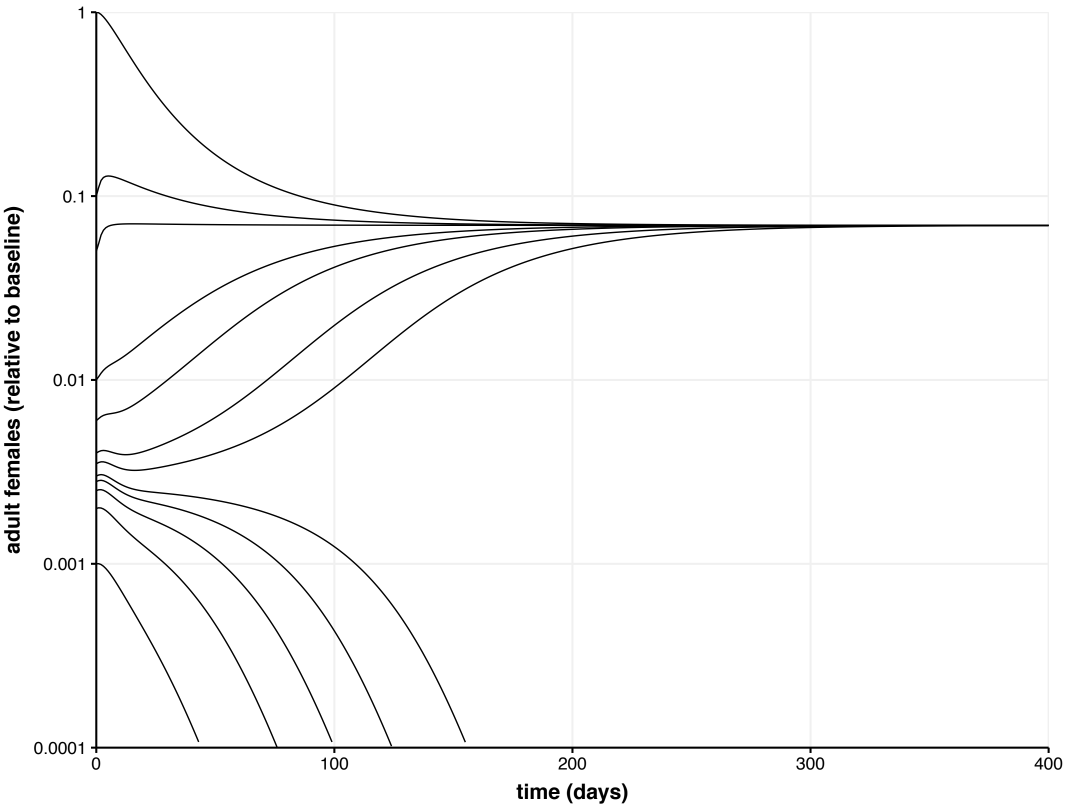


**Figure S2: Bistability of a 1-locus LFK system with release below $r_c$.** Each line plots the trajectory of number of females (relative to the wild-type equilibrium) over time with a release ratio of $r = 2$ beginning at time 0. A log scale for the number of adult females is used in order to illustrate behavior at small values. The initial number of adult males and juveniles are reduced by the same amount as the females (0.01 begins with 1/100 of the wild-type equilibrium count of each class). Simulations starting from low counts of wild-type result in population extinction, whereas higher initial counts result in the population reaching a non-zero equilibrium. Note that only a single dimension of the system is plotted, which means the time-series can exhibit different dynamics at the same value of relative adult females. Fitness parameters are equal to the default values from the main text: $s^H = 0.2$, $c_A = 0.55$, and $s^M = 0.1$.

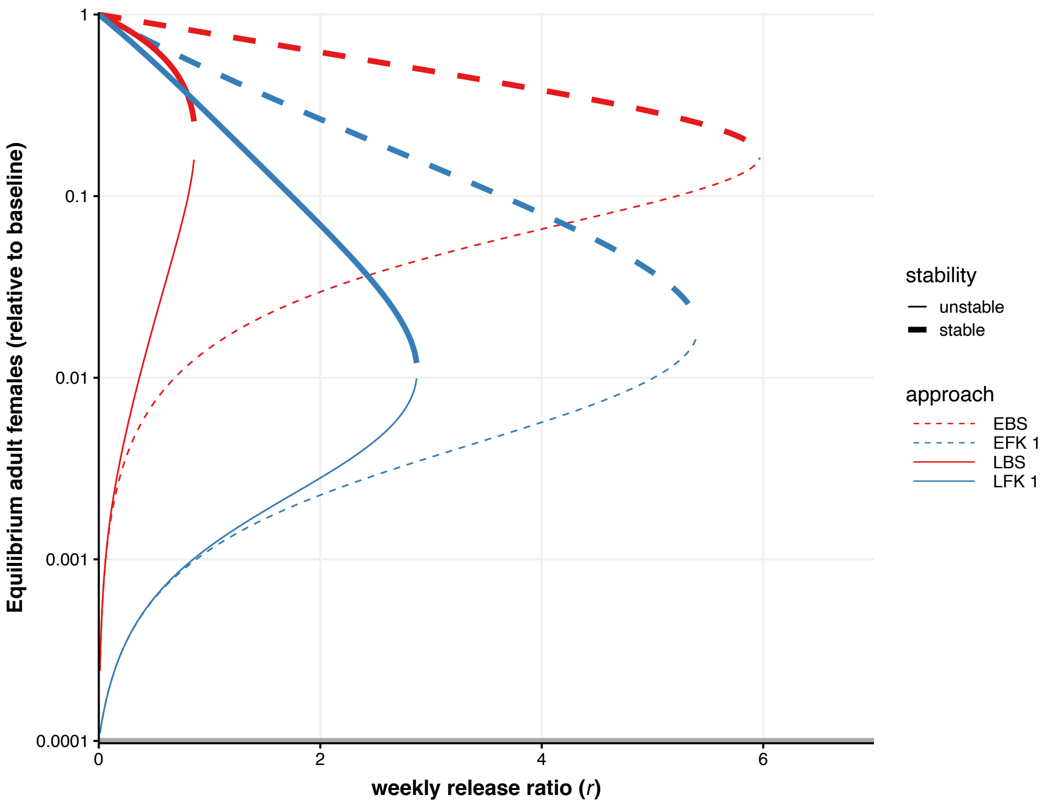


**Figure S3: Bifurcation diagram for 1-locus approaches, with the default parameter values from the main text.** Analytical solutions and their corresponding eigenvalues were calculated. Here, the number of adult females in real, positive solutions is plotted for varying $r$. A log scale for the number of adult females is used in order to illustrate small equilibria. The grey line at the x-axis indicates the stable solution of 0 for all approaches (an extinct population with only released adult males remaining). Otherwise, approach varies with line color and type as in previous figures. The systems exhibit a saddle-node bifurcation, where at small values of $r$, there are two stable solutions (thick lines, one at zero and one positive) with an unstable (thin lines) solution in between. When $r$ reaches $r_c$, the positive stable and unstable solutions collide. For larger $r$, the only equilibrium is the stable equilibrium at 0.

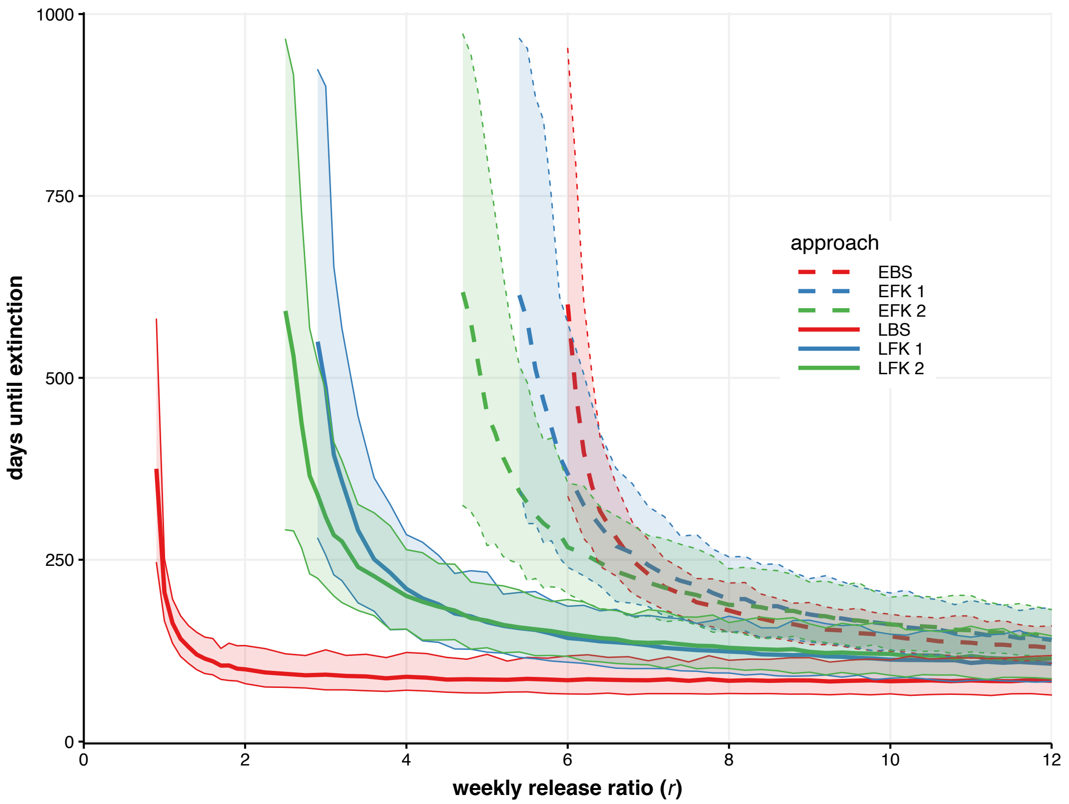


**Figure S4: Time until no adult females remaining in stochastic simulations.** For each approach and value of *r*, 300 stochastic simulations were conducted for up to 1000 days. Thick lines show the mean time until there were zero adult females left in the population, and the ribbons show the 2.5% and 97.5% quantiles. Data is only plotted for values of *r* where at least 200 simulations resulted in 0 adult females by day 1000. Color and line type indicate approaches as in previous figures. Parameters are the default in the main text: $\beta = 3$, $s^H = 0.2$, $s^M = 0.1$, $c_A = 0.55$, and $h = 0.5$. The mean times are similar to Figure 2B in the main text, which shows that the time until falling below 1 adult female (from 2000 wild-type at equilibrium) in deterministic simulations is a good approximation of the stochastic simulations.

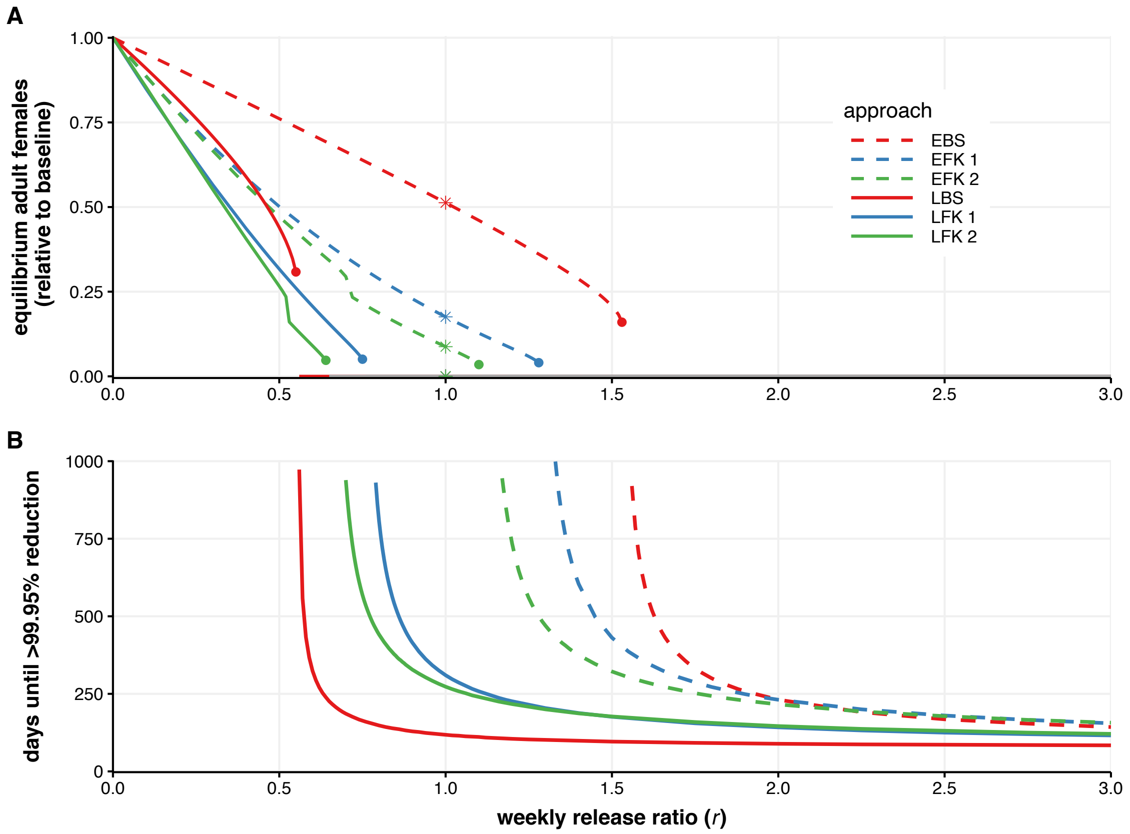


**Figure S5: Release outcomes across different release ratios with β = 2. A:** Long-term, stable equilibria for number of viable adult females (relative to equilibrium) for different *r*, found by simulating the system of differential equations until at steady-state. The asterisks indicate $r = 1$, for which the equilibrium values correspond to the simulations in the left-column, center-row panel of Figure S1. Each genetic approach exhibits a bifurcation at a critical release ratio, $r_c$, above which extinction is guaranteed. Above $r = 0.64$, two or more approaches lead to extinction of the population and hence have equilibria at zero: this is indicated using a grey line. Each of the 2-locus FK (green lines) approaches exhibit a discontinuity where *r* becomes large enough to drive one of the alleles to fixation, decreasing the equilibrium population size (as explained in Appendix 3 and illustrated in Figure S6 and S7). **B:** Time until the number of viable adult females is under 0.05% of equilibrium in deterministic simulations for different *r*. Color and line type match that of previous figures. Fitness parameters are equal to the default values from the main text: $s^H = 0.2$, $c_A = 0.55$, and $s^M = 0.1$.

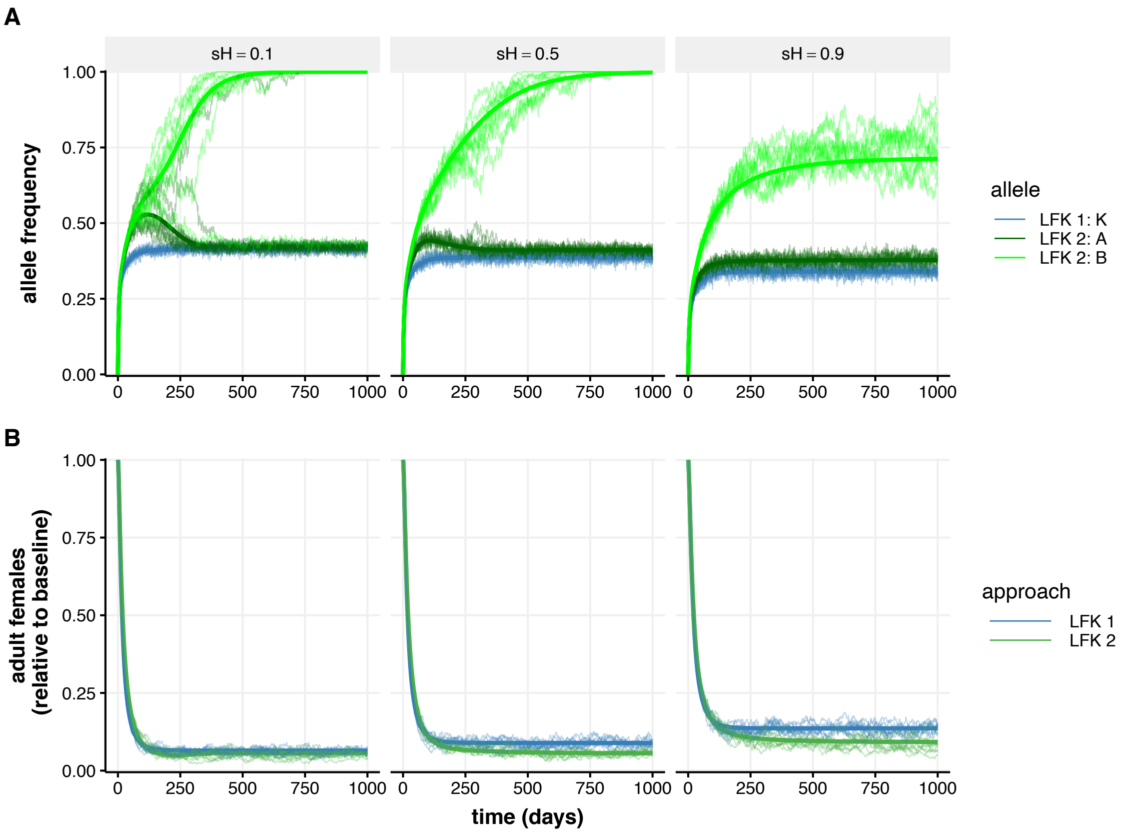


**Figure S6: 2-locus LFK juvenile allele frequencies over time at $r = 2$ for different hatching fitness costs. A:** Allele frequency in juveniles shows different possible outcomes of A (dark green) and B (light green) allele frequencies for 2-locus LFK, and the K allele frequency in 1-locus LFK. The maximum possible allele frequency for 1-locus (or 2-locus when one allele is at fixation) is 0.5, when all individuals are heterozygous. **B:** Corresponding relative adult females in the population over time. Fitness parameters are $c_A = 0.75$, $s^M = 0.25$, and $h = 0.25$. Deterministic simulations (thick lines) and 10 stochastic simulations (thin lines) are shown. At low hatching fitness costs (left column), the B allele goes to fixation in most simulations, but the A allele can also go to fixation despite having higher costs than B. At moderate costs (middle column), the B allele always goes to fixation, and at high costs (right), both alleles reach intermediate frequencies. See Figure S7 for the outcomes of allele frequencies across parameter space.

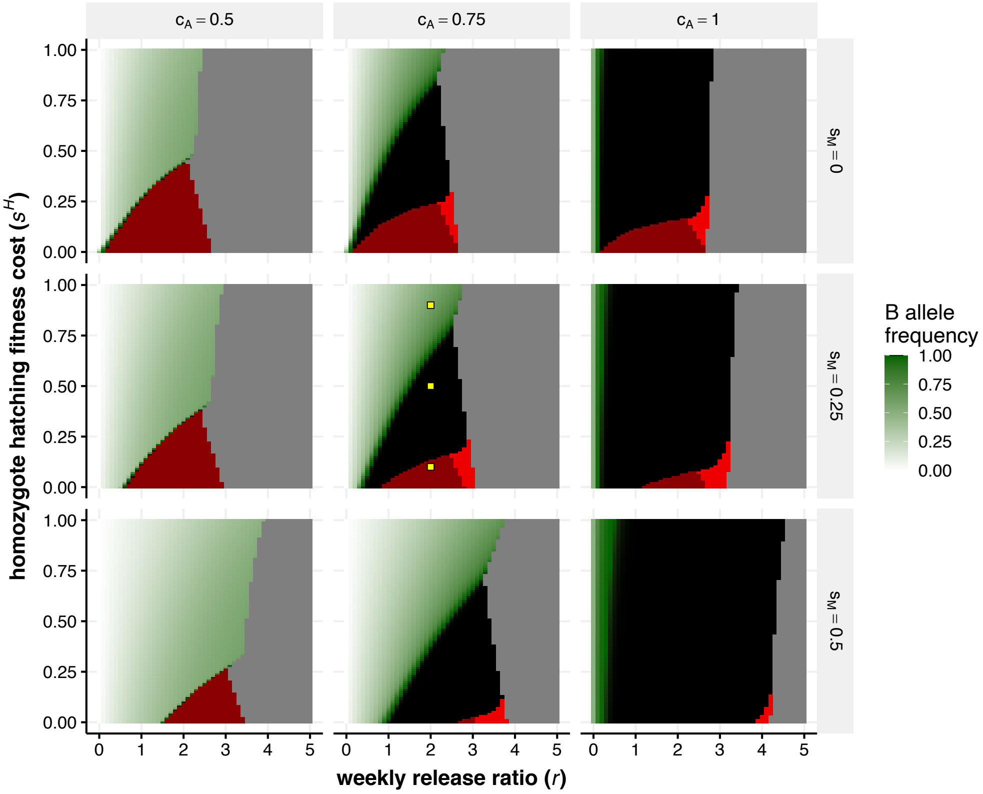


**Figure S7: Allele frequency outcomes from deterministic simulations of 2-locus LFK for various fitness parameters and release ratios when $h$ = 0.5**. At each unique set of fitness parameters ($s^M$ differing between rows of panels, $c_A$ differing between columns of panels, and $s^H$ differing across the y-axis of each panel) and weekly release ratio ($r$, differing across the x-axis of each panel), long-term outcomes of two simulations with different initial conditions were compared. The systems were perturbed by adding transgenic adult male and females to the system at time 0. One simulation began with aaBB and the other AAbb males and females in equal numbers to the wild-type equilibrium number of males and females. In some regions of parameter space, $r$ is large enough (toward the right side of each panel) to cause the system to go extinct in both simulations (grey regions). At lower $r$, there are four potential outcomes, three of which are illustrated in Figure S6 (panel parameters indicated by yellow points): 1. the A and B alleles end at an intermediate frequency in both simulations (B frequency is indicated by shade of green), 2. the B allele goes to fixation in both simulations (black regions), 3. whichever allele was at higher frequency at time 0 goes to fixation (dark red), or 4. the B allele goes to fixation if starting at a higher frequency, while the system goes extinct if beginning with additional A alleles in the population (light red).
